# Controlled delivery of immunomodulatory factors for mineralized tissue formation in an inflammatory microenvironment

**DOI:** 10.1101/2025.05.21.655332

**Authors:** Katherine H. Griffin, Isabel S. Sagheb, Thomas P. Coonan, Langston A. Wu, Douglas J. Rowland, Boaz Arzi, Jamal S. Lewis, J. Kent Leach

## Abstract

Mesenchymal stromal cells (MSCs) are a promising cell-based therapy for bone healing, contributing to tissue regeneration through direct differentiation or immunomodulatory factor secretion. However, diseases that feature chronic or dysregulated inflammation, such as non-union fractures and osteonecrosis of the jaw (ONJ), have proven difficult to treat with current MSC-based approaches. Here, we investigated whether controlled delivery of immunomodulatory factors would allow MSCs to simultaneously undergo osteogenic differentiation and modulate inflammation. We first used a Design of Experiments approach to identify the type and concentrations of immunomodulatory factors (IMFs) that most effectively induce concurrent pro-regenerative macrophages and MSC osteogenic differentiation, then loaded these IMFs into polymeric microparticles for controlled release. Through our *in vitro* models, we demonstrated that microparticles releasing IL-10 and IL-4 promote naïve MSC osteogenesis and modulate immune response, even in chronic, physiologically relevant, inflammatory conditions. We then applied this approach to an *in vivo* rat model of ONJ as a clinically relevant example of such conditions. We observed clinically relevant sex-based differences in inflammation and bone formation that have not yet been reported. These data represent key findings that will facilitate the reversal of diseases that are linked to chronic bone loss and inflammation, such as ONJ.

## 1. Introduction

Mesenchymal stromal cells (MSCs) are a promising cell-based therapy for bone regeneration and have been widely explored to treat musculoskeletal disease. For over a decade, systemic or local MSC injection was attempted as an experimental therapeutic for muscle, bone, and cartilage regeneration [1, 2]. However, success was highly variable and often limited by poor cell survival [3, 4]. Biomaterials can augment MSC-mediated tissue repair through direct differentiation cues or by promoting trophic factor secretion to recruit endogenous cells and modulating the local immune environment [5, 6]. The latter is defined as the MSC secretome, and it has been widely used as a cell-free approach to elicit many of the same effects as MSCs [7, 8]. A few studies have investigated direct differentiation contributions of various stromal cell therapies delivered in hydroxyapatite [9, 10], as local platelet-rich plasma injections [11], and as cellular sheets [12], with general improvements in bone regeneration. However, these studies largely fail to leverage the benefits of the immunomodulatory MSC secretome and its subsequent effects on long-term bone formation. Harnessing both regenerative processes of MSCs with concurrent differentiation and secretome production offers increased therapeutic potential yet has been difficult to achieve.

The role of the immune system is an understudied element of many musculoskeletal conditions, especially those characterized by increased or dysregulated inflammation [13]. This has thwarted therapeutic advancement for numerous pathologies, from primary idiopathic conditions like non-union fractures [14] to induced sequelae such as medication-related osteonecrosis of the jaw (MRONJ) [15] and even autoimmune disorders like rheumatoid arthritis [16]. For years, this has been addressed clinically though combinations of systemic antibiotics, non-steroidal anti-inflammatory (NSAID) administration, or immunomodulation with stronger biologics, such as tumor necrosis factor (TNF) blockers [17, 18]. Additional approaches to correct the altered regulation of the inflammatory environment are warranted, as these treatments are not specific to disease pathophysiology and hold serious complications related to antimicrobial stewardship and off-target effects. Some studies have explored the immunomodulatory potential of MSC-mediated treatment [2], with MSC conditioned media [19, 20] or exosomes [21, 22]. Yet these systemic, short-acting strategies fail to provide targeted, sustained regenerative cues that interrupt the inflammatory positive feedback loops that are distinct and critical in these chronic diseases. Given these harsh microenvironments, additional cues may be necessary to both resolve inflammation and promote bone regeneration. While MSCs upregulate bioactive factor production in order to modulate the inflammatory environment [23], it is unknown if the controlled delivery of similar biomolecules may allow MSCs to persist longer and undergo osteogenic differentiation toward the osteoblastic phenotype.

Here, we locally deliver immunomodulatory factors and interrogate MSC differentiation in physiologically relevant inflammatory conditions, such as those found in non-union fractures, MRONJ, and rheumatoid arthritis. We hypothesize that the local delivery of these immunomodulatory factors (IMFs) will decrease the pro-inflammatory microenvironment *via* altered macrophage polarization, thereby indirectly promoting the osteogenic potential of MSCs. We use a Design of Experiments approach to optimize the IMF formula for maximal osteogenesis and immunomodulation, load IMFs into polymeric microparticles for controlled release, characterize cellular response to these IMF microparticles within hydrogels, and challenge this treatment strategy in an *in vivo* rat model of MRONJ.

## 2. Methods

### Cell culture

Bone marrow-derived Sprague Dawley rat MSCs (rMSCs, Cyagen Biosciences, Santa Clara, CA) were cultured in αMEM (Invitrogen, Carlsbad, CA) supplemented with 10% fetal bovine serum (FBS) (Biotechne, Minneapolis, MN) and 1% penicillin-streptomycin (P/S) (Gemini Bio Products, West Sacramento, CA) in standard conditions (37°C, 21% O_2_, 5% CO_2_). rMSCs were used at passage 4-5 for all experiments. For osteogenic studies, rMSCs were maintained for 21 days in either basal media for naïve rMSC experiments or osteogenic media (OM) consisting of α-MEM supplemented with 50 μg/mL ascorbate-2-phosphate and 10 mM β-glycerophosphate (both from Sigma-Aldrich, St. Louis, MO). Dexamethasone was not included, since a major aim of this study is to evaluate the immune response, which is greatly altered with corticosteroids.

IC-21 murine macrophages (ATCC, Manassas, VA) were cultured in RPMI 1640, ATCC Modification (Gibco, Thermo Fisher, Waltham, MA) supplemented with 10% FBS and 1% P/S. Macrophages were used at passage 6-10 for all experiments and maintained below 70% confluency.

### Design of Experiments (DOE) to determine effect of IMFs on osteogenesis and macrophage polarization

We used a Response Surface Methodology Design, specifically Box Behnken [24, 25], to determine the concentrations of IL-4, IL-10, and prostaglandin E_2_ (PGE_2)_ that concurrently maximize macrophage polarization and rMSC osteogenesis. We generated 13 unique experimental conditions with Design-Expert 11 (Stat-Ease, Minneapolis, MN), in which we treated IC-21 macrophages for 3 days and rMSCs for 21 days in OM with 0-100 ng/mL IL-4, 0-100 ng/mL IL-10, and 0-2 μg/mL PGE_2_ (**Supplemental Tables 1** and **2**). These concentration ranges were determined through compiled data from the literature for common and physiologically relevant *in vitro* treatments, as well as a pilot dosing study (*data not shown*). Resultant macrophage polarization was analyzed with flow cytometry, and osteogenic potential was measured with alkaline phosphatase activity (ALP), Alizarin red staining, and calcium quantification. The predicted significance and interactions of IMF treatments on these responses were determined with the surface response curves generated by the Design-Expert Software (**Supplemental Figure S1, Figure 1**).

**Figure 1:**
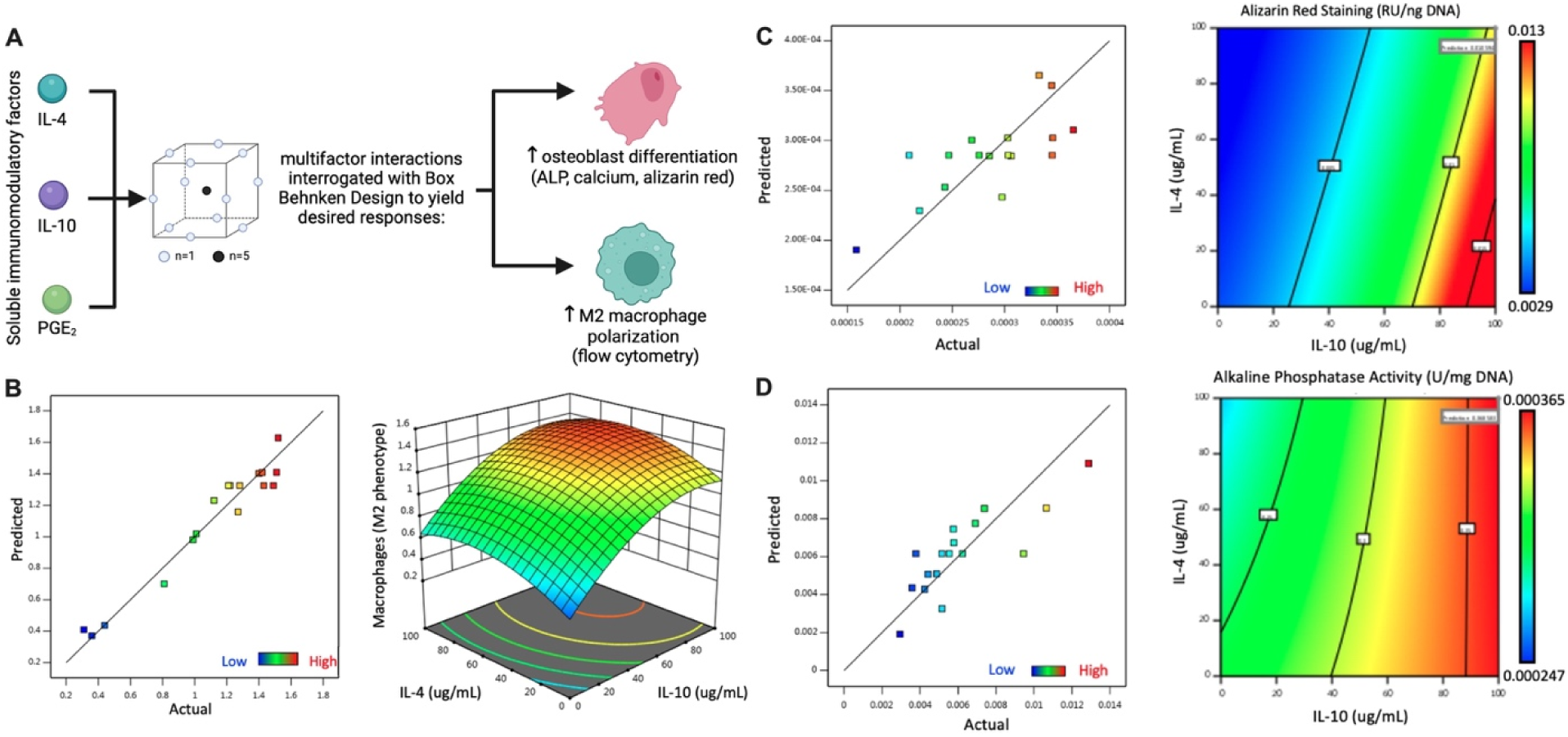
Combined presentation of IL-4 and IL-10 maximize anti-inflammatory macrophage polarization and rMSC osteogenesis based on DOE approach. (A) Schematic of DOE conditions with input factors, specific design, and output responses. Predicted vs. Actual plots [35] (left) and surface response plots (right) for (B) macrophage polarization, (C) AR staining, and (D) ALP activity.

### Biochemical characterization of osteogenic differentiation

The biochemical outputs were normalized to total DNA content, which was quantified with Quant-iT PicoGreen dsDNA Assay Kit (Invitrogen, Carlsbad, CA) according to the manufacturer’s protocol. Other osteogenic studies showed no differences in DNA content (*data not shown*) and as such were presented without normalization. Intracellular alkaline phosphatase (ALP) expression was quantitatively assessed from passive lysis buffer lysate using a routine p-nitrophenyl phosphate (PNPP) colorimetric assay wherein absorbance is measured at 405 nm. Calcium deposition was quantified using a Stanbio™ Calcium Liquid Reagent kit (Thermo Fisher, Chicago, IL) according to manufacturer’s instructions after digestion in 1M HCl for 24-48 h. Alizarin red (AR) staining was used to assess calcification. Samples were fixed in 10% buffered formalin for either 30 min at room temperature or overnight at 4°C, rinsed twice with deionized (DI) water, and incubated with 1% (w/v) Alizarin red (Sigma) in DI water for 2 min. Samples were then washed with DI water until no visible background color remained in internal control blank wells. For the DOE, AR was quantified with stain extraction using 10% (w/v) cetylpyridinium chloride monohydrate (Sigma) in DI water for 2 h, after which absorbance was measured at 540 nm.

### Flow cytometry for macrophage polarization

For all polarization experiments, polarization controls in monolayer were treated with basal media (M0), 100 ng/mL LPS (M1), and 40 ng/mL IL-4 (M2) (*data not shown*). Monolayer cells were collected with ice-cold 2.5 mm EDTA in PBS and gentle scraping. Samples in 10% GelMA were digested in 1.5 mg/mL collagenase D in 3% FBS in PBS at 37°C for 30 minutes to fully degrade the gels. Cell suspensions were filtered through 70 μm filters, spun down, and resuspended at 37 °C 3% FBS in PBS. Then, cells were stained for flow cytometry as previously described [26, 27]. Following Fc_y_ receptor blocking (1:40, TruStain FcX, BioLegend), cells were stained with antibodies against F4/80 (1:50, eBioscience, cat. MF48021), CD86 (1:160, eBioscience, cat. 47-0862-82), and CD206 (1:40, eBioscience, cat. 48-2061-82). Cellular viability was evaluated with fixable Zombie Aqua (1:250, LifeTech). Cells were fixed with PFA (2%), permeabilized with Triton-X (0.1%), and stained for intracellular markers iNOS (1:500, eBioscience, cat. 12-5920-82) and Arginase-1 (1:500, eBioscience, cat. 53-3697-82) overnight at 4°C with gentle agitation. Macrophages with an M1 phenotype were characterized by F4/80+CD86+iNOS+ populations and M2 phenotypes by F4/80+CD86-CD206+ARG1+ populations.

For experiments investigating macrophage response to rMSC conditioned media, rMSC media was collected under sterile conditions after incubation with rMSCs for 24 h and stored at −80°C until use. IC-21 macrophages were treated with conditioned media at a 1:1 ratio with basal media and incubated for 24 h. Media was then replaced with basal media, cells were incubated another 24 h, then collected for flow cytometry. Polarization controls for these experiments were also mixed with 1:1 basal macrophage media and basal αMEM to control for confounding effects from added αMEM.

### Microparticle synthesis and characterization

Microparticles were formed using an established double emulsion technique [28, 29] using a 50:50 polymer composition of poly(lactide-co-glycolide) (PLGA; PURASORB PDLG 5004A, Corbion, Netherlands) in methylene chloride (MeCl; Sigma). Polyvinyl alcohol (PVA; MW ∼15,000 g/mol; MP Biomedicals, Solon, OH) was used as an emulsion stabilizer. Briefly, 250 mg of PLGA was dissolved in MeCl for 15 min at 150 RPM at 20% w/v ratio. Recombinant human IL-4 or IL-10 (Cat# 200-04 and Cat# 200-10, respectively, Peprotech, Cranbury, NJ) were reconstituted according to manufacturer’s instructions at 0.8 mg/mL in DI water or 5 mM sodium phosphate pH 7.2, respectively. Blank microparticles were loaded with PBS. These aqueous solutions were individually added at a 1:10 ratio to the PLGA-MeCl solution and emulsified for 60 s. The primary emulsion was quickly added to 2.5 mL of 3% PVA in DI water and vortexed again for 60 s to form the secondary emulsion. This solution was added dropwise to 75 mL of 1% PVA solution on a magnetic stirrer, covered with a lab wipe, and left stirring overnight to evaporate residual MeCl. Microparticles were collected and washed 3 times with DI water and centrifugation at 8,000 x g for 15 min per wash. Following the final wash, microparticles were frozen at −80°C then lyophilized for 24-48 h. Microparticles were stored at −20 °C until used.

Following the final wash, a small aliquot of particles was collected for size characterization with S3500 Laser Diffraction Analyzer (Microtrac, Verder Scientific, Newton, PA), as well as morphological characterization *via* scanning electron microscopy (SEM). After lyophilization, we conducted 10-day release kinetic studies, with 1 mL PBS collected at each timepoint such that the total solutions collected over time amassed the total secreted analyte. At each collection, samples were centrifugation at 8,000 x g for 5 min, supernatant was collected, and microparticles were resuspended in 1 mL PBS. Results were quantified as determined by Quantikine ELISA Kits specific for each IMF (R&D Systems, Minneapolis, MN) and then normalized to the initial known starting mass of each replicate and theoretical total amount of analyte.

### Fabrication of gelatin methacryloyl hydrogels

Gelatin methacryloyl (GelMA; 300 bloom, MilliporeSigma, St. Louis, MO) was dissolved at 20% (w/v) in αMEM with 0.6% (w/v) 2-Hydroxy-4′-(2-hydroxyethoxy)-2-methylpropiophenone (Irgacure 2959 in αMEM) with agitation at 80°C for 30Umin protected from light. Lyophilized microparticles were suspended in either αMEM or cell suspensions (1.0×10^6^ for *in vitro* studies, 3.0×10^7^ for *in vivo* studies) and resultant solutions were mixed with an equal volume of the GelMA to yield a final concentration of 10% GelMA, 0.3% Irgacure, 15 mg/mL total microparticles, and 5.0×10^5^ cells/mL for *in vitro* studies or 1.5×10^7^ cells/mL for *in vivo* studies. The final solution was cast into 8 mm diameter x 1 mm high molds (75 μL) for *in vitro* studies and 4-mm-diameterU×U1-mm-high molds (20 μL) for *in vivo* studies. Constructs were exposed to 17UmW/cm^2^ ultraviolet (UV, 320-500 nm) light for 2Umin before culture for *in vitro* experiments or immediate *in vivo* use.

### Mechanical characterization

The shear storage moduli of 8 mm diameter gels were determined using a Discovery HR2 Hybrid Rheometer (TA Instruments, New Castle, DE) with a stainless steel, cross hatched, 8 mm plate geometry. An oscillatory strain sweep ranging from 0.004% to 4% strain with an initial axial force of 0.2 N was performed on each gel to obtain the linear viscoelastic region (LVR) before failure [30]. Gel shear storage modulus was calculated from the average of at least seven LVR data points.

### Quantitative polymerase chain reaction (qPCR)

PCR samples were collected in in 300 μL TRIzol (Invitrogen), from which total RNA was isolated as per the manufacturer’s instructions. Samples were disassociated *via* sonication at 40% amplitude in 10 s increments on ice until achieving a homogenous solution. 800 ng of RNA was reverse transcribed to complimentary DNA using the QuantiTect Reverse Transcription Kit (Qiagen, Germantown, MD) and normalized to a final concentration of 20Ung/μL.

qPCR was performed using Taq PCR Master Mix (Qiagen) in a QuantStudio 5 real-time PCR system (Thermo Fisher). Rat specific primers (all from Thermo) for Gapdh (Hs02786624_g1), Rpl13 (Rn00821947_g1), Runx2 (Rn0101512298_m1), and Col1A1 (Rn01463848_m1) were used to analyze gene expression. All genes were normalized to the endogenous control housekeeping genes, either *Gapdh* or *Rpl13*, to yield a ΔCt value. Gene expression was further normalized to the appropriate negative control in each experiment to determine the ΔΔCt. Fold change was calculated using the 2^−ΔΔCt^ method.

### Quantification of cytokine production

Immediately before sample collection for PCR, media conditioned for 24 h was collected for multiplex proteomic analysis and frozen at −80 °C until use. Samples were prepared for a ProcartaPlex Rat Cytokine & Chemokine Panel 22plex (Lot#: 336326-001, Cat. EPX220-30122-901; Thermo Fisher Scientific) per manufacturer’s instructions, incubated at 4°C overnight, processed on a Luminex lx200 (Luminex Corp, Austin, TX), and analyzed with ThermoFisher Connect ProcartaPlex Analysis App. For each analyte, readings beyond the upper limit of detection were reported as the highest pg/mL concentration on the standard curve, and readings below the lower limit of detection were reported as 0 pg/mL. Results were then normalized to a percentile scale as described in the figure legends.

### In vivo model of MRONJ

Animals were treated in accordance with all University of California, Davis animal care guidelines and National Institutes of Health (NIH) animal handling procedures under IACUC protocol #23471. 8-week-old male and female Sprague Dawley rats (NTac:SD; females: 175-225 g, males: 225-290 g, Taconic, La Jolla, CA) were intravenously injected with USP pharmaceutical grade zoledronic acid (Sigma) in PBS at 0.1 mg/kg 3 weeks prior to and immediately following tooth extraction [31]. For the surgical procedure, animals were anesthetized and maintained under a 1–3% isoflurane/O_2_ mixture delivered through a nose cone. Prior to tooth extraction, animals were placed in supine recumbency, and a local infiltration block was made in the gingiva immediately adjacent to the right maxillary first molar tooth using 50 μg bupivacaine. A small gauze was placed in the oropharynx to prevent aspiration of fluids. A Schmidtke small mouth gag and a small pouch dilator (speculum) were placed to facilitate an intraoral approach, and the right maxillary first molar was extracted using a Crossley rabbit molar luxator. Alveoloplasty was performed using a round diamond bur on a high-speed handle piece with constant saline irrigation to the alveolar bone. The empty alveolus bone defects were immediately packed with GelMA gels containing rMSCs, IMF microparticles, or both rMSCs and IMF microparticles. The surgical site was closed with 5-0 Monocryl suture in a simple interrupted fashion (Y463, Ethicon, Raritan, NJ). Buprenorphine (0.05 mg/kg) was administered twice per day for two days as analgesia. To avoid excess pain and confounding trauma to the surgical site, rats were fed moist chow for 7 days.

### Preclinical imaging of local inflammation

Immediately following surgery, animals were intravenously injected with 300 μL IVISense Pan Cathepsin 750 (NEV10001EX, Revvity, Hopkinton, MA). Twenty-four h after injection, fluorescent signal was detected at 750 nm using an In Vivo Imaging System (IVIS) Spectrum (PerkinElmer, Waltham, MA) with animals in left lateral recumbency. Images were processed with ROI selection and quantified using the IVIS’s accompanying software to quantify the presence of inflammation in the maxilla. Quantifications solely include male specimens, as female rats had significant background signal around the eye that interfered with the molar ROI.

### Microcomputed tomography (microCT)

Prior to surgery (baseline), within 24 h after surgery (D0), and six weeks after surgery (W6), all animals designated for microcomputed tomography (microCT) underwent live-animal microCT imaging. Animals were anesthetized and maintained under a 1–3% isoflurane/O_2_ mixture delivered through a nose cone, then imaged (80 kVp, 150 μA) at 50 μm resolution using GNext PET/CT (Xodus Imaging, Torrance, CA) A phantom probe with known hydroxyapatite densities was also scanned with the same protocol. Reconstructed images were then analyzed in Dragonfly 3D (version: 2022.2.0.1409, Comet Technologies, San Jose, CA). Volumes of interest (VOIs) were drawn manually plane-by-plane over the right maxillary first molar and associated maxilla of the first baseline scans. Subsequent scans were then registered to the baseline scan for quantification using this VOI. Registration of post extraction scans to the baseline were performed manually to along the skull and upper maxilla. Bone volume was quantified as the voxels with hydroxyapatite density greater than 250 mg/mL. Mineral density was interpolated from a standard curve generated from the known phantom densities.

### Histology

Three days after surgery and one day after W6 microCT scans were performed, rats were euthanized, and maxilla were isolated and fixed in 10% buffered formalin at 4°C for 24-48 h. Samples were rinsed twice with PBS, demineralized in Calci-Clear (National Diagnostics, Atlanta, GA) for 2 weeks with solution changes every 2-3 days, then fixed again in 10% buffered formalin at 4°C for 24-48 h. Samples underwent tissue processing, paraffin-embedding, and sectioning at 10 μm thickness.

H&E staining was performed as previously described, and slides were imaged with 4X and 10X objectives using a Nikon Eclipse TE2000U microscope. For immunohistochemistry staining, slides were rehydrated and exposed to heat mediated antigen retrieval with a sodium citrate buffer for 20 min. Samples were then incubated in blocking buffer composed of 10% goat serum and 10 mg/mL Bovine Serum Albumin (BSA) for 30 min at room temperature. For granulocyte detection, slides were incubated with recombinant anti-granulocytes monoclonal antibody (ab33760, Abcam) at a concentration of 1:100 for 1 h at room temperature. Slides were then treated with a secondary goat anti-mouse antibody conjugated to Alexa fluor 594 (ab150118, Abcam) at a concentration of 1:250 for 1 h at room temperature. For angiogenesis, slides were incubated with recombinant anti-CD31 antibody (ab182981, Abcam) at 1:200 for 1 h at room temperature, then treated with a secondary goat anti-rabbit antibody conjugated to Alexa fluor 488 (ab150081, Abcam) at a concentration of 1:300 for CD31 for 1 h at room temperature. Slides were counterstained with DAPI (Thermo Fisher Scientific). Slides were imaged with a confocal microscope (Leica Stellaris 5, Leica Microsystems, Deerfield, IL). Positive signal was quantified with ImageJ. Quantifications were determined from 3 fields of view on at least 3 different slides per group by a blinded observer.

### Statistical analysis

Data are presented as mean ± standard deviation for *in vitro* data and mean ± standard error of the mean for *in vivo* data. Unless otherwise stated, *in vitro* data represent technical replicates. Statistical analysis was conducted with Prism 10.0.0 (GraphPad, San Diego, CA) software utilizing either unpaired t-test assuming Gaussian distribution, one-way or two-way analysis of variance (ANOVA) with post hoc Tukey’s test depending on the number of groups and comparisons or paired two-way ANOVA with post hoc Tukey’s test for mineral density data. We reported Spearman’s correlation coefficients, and associated significance was tested with non-parametric, two-tailed t-tests with 95% confidence intervals. F-tests to compare variances were also conducted. Groups with different letters indicate statistically significant differences (p < 0.05), while groups with the same letters were not significant.

## 3. Results

We first examined whether a chronic, physiologically relevant, inflammatory environment was inhibitory to osteogenesis. Acute levels of TNFα-induced inflammation are necessary for MSC-mediated osteogenesis in clinical fracture patients, and corroborating *in vitro* studies show similar trends though typically treating on the order of nanograms [32, 33]. However, these levels are supraphysiologic compared to those found with the chronic inflammation, which are reported on the order of picograms [34, 35]. Thus, we cultured rat MSCs (rMSCs) in osteogenic media supplemented with 50 pg/mL TNFα media transiently for 3 days or constantly for 21 days to understand the effects of low-level inflammation on osteogenic potential (**Supplemental Figure S1A**). We found that neither transient nor constant TNFα affected rMSC proliferation (**Supplemental Figure S1B**) or osteogenesis as characterized by Alizarin Red (AR) and alkaline phosphatase (ALP) activity (**Supplemental Figure S1C, S1D**).

To halt the inflammatory signals and promote tissue regeneration, we next investigated a combination of immunomodulatory factors (IMFs) to promote an anti-inflammatory, pro-regenerative macrophage phenotype and instigate osteogenesis. Classically referred to as M2 macrophages, anti-inflammatory macrophages are further sorted into subsets based on activating cytokines and functional responses [36]. Generally, IL-4, IL-10, and Prostaglandin E_2_ (PGE_2_) are known to induce the two most prevalent M2 subsets, M2a (IL-4) [37] and M2c (IL-10 and PGE_2_) [37, 38] macrophages, respectively. Therefore, we used a Design of Experiments (DOE) approach to define the combination of IL-4, IL-10, and PGE_2_ that would maximize M2 macrophage polarization and rMSC osteogenesis. Using a Box Behnken Design, we treated rMSCs for 21 days and IC-21 macrophages for 3 days, with 13 unique combinations of IL-4, IL-10, and PGE_2_ (**Supplemental Tables 1, 2**). Macrophage polarization was evaluated with flow cytometry (M2 defined as CD86-CD206+ARG1+), and rMSC differentiation was evaluated with AR staining, ALP activity, and calcium deposition (**Supplemental Figure S2**). Predicted vs. Actual graphs were closely aligned for macrophage response (**Figure 1B**), AR staining (**Figure 1C**), and ALP activity (**Figure 1D**). Surface response plots (**Figure 1B-D**) showed that treatment of macrophages and rMSCs with 100 ng/mL IL-4, 100 ng/mL IL-10, and 0 μg/mL PGE_2_ maximized M2 polarization and osteogenic differentiation.

Immunomodulatory factors (IMFs) IL-4 and IL-10 were next loaded into unphagocytosable PLGA microparticles for controlled release of 100 ng/mL of each factor. Size characterizations of IL-4 (**Figure 2A, Supplemental Figure S3**) and IL-10 (**Figure 2C, Supplemental Figure S4**) microparticles revealed normally distributed diameter measurements with an average of 55.1 ± 27.9 μm and 65.0 ± 24.0 μm, respectively. Both IMF microparticles exhibited similar, generally spherical, morphology *via* scanning electron microscopy (**Figure 2B, 2D**), and release studies demonstrate a classic burst release kinetic with near 100% IMF delivery by day 3 (**Figure 2E**).

**Figure 2:**
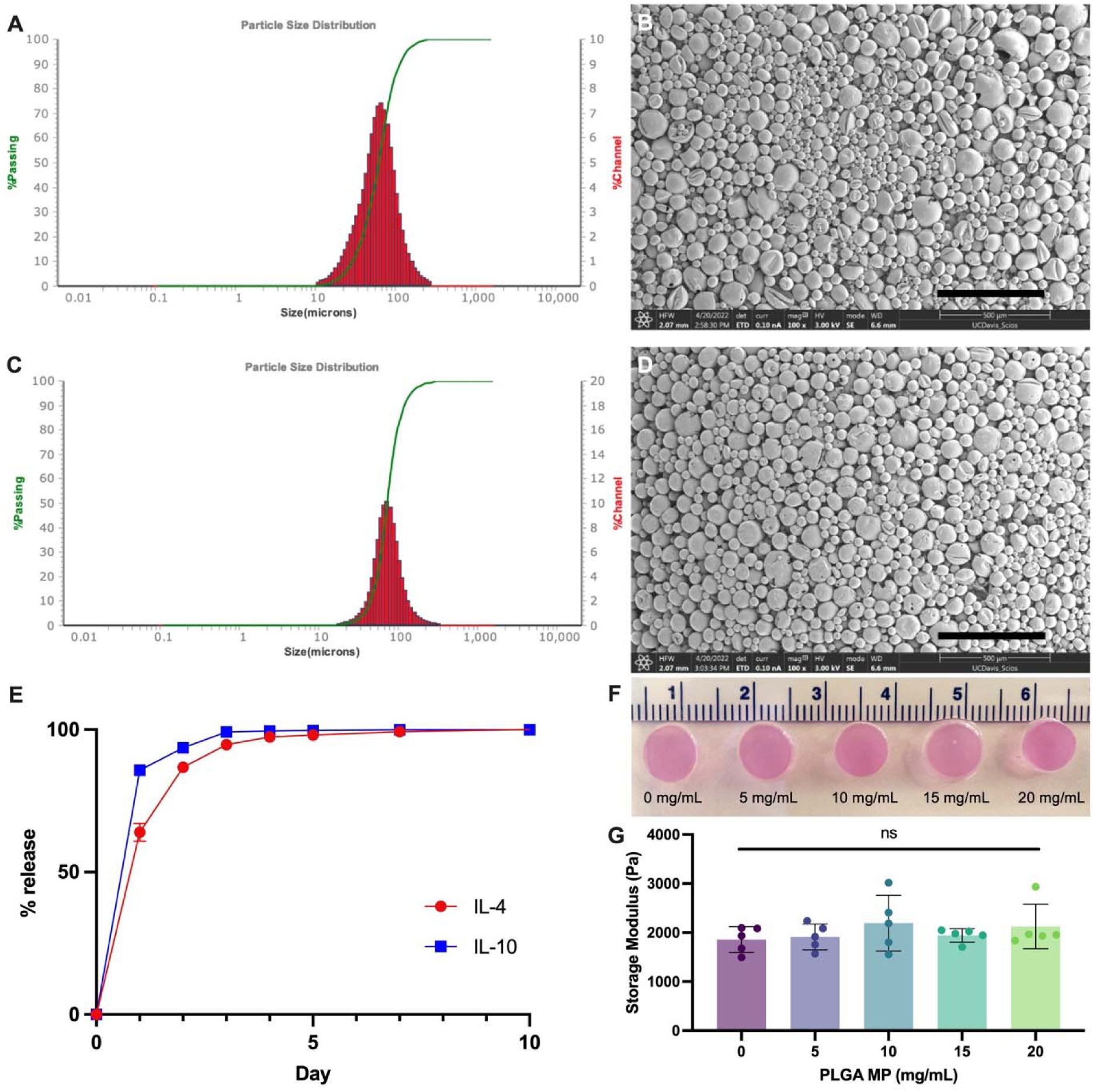
PLGA microparticles loaded with IL-4 and IL-10 exhibit similar size, morphology, and release kinetics, and can be loaded into 10% GelMA with no changes in mechanical properties. Size distribution graphs and representative SEM image of IL-4 **(A, B respectively)** and IL-10 **(C, D respectively)** microparticles. Scale bar = 500 μm. **(E)** Release kinetics over 10 days as measured by protein-specific ELISA, data are mean ± SD (n=4). **(F)** Gross images of GelMA hydrogels loaded with microparticles and **(G)** shear storage modulus of 10% GelMA hydrogels loaded with 0-20 mg/mL MPs. Numerical demarcations on gross images = 1 cm. Data are mean ± SD (n=5). ns denotes no significance based on on-way ANOVA.

To facilitate targeted, localized delivery, microparticles were loaded into 10% GelMA hydrogels. With an eye towards *in vivo* applications, we required hydrogels to maintain integrity under load and measured the effect of microparticle incorporation on hydrogel storage modulus. To conserve growth factors, blank PLGA microparticles of comparable size (average diameter = 44.8 μm) and morphology (**Supplemental Figure S5**) were used for these experiments. We observed no change in gross hydrogel morphology (**Figure 2F**) or shear storage modulus (**Figure 2G**) when loaded with 0-20 mg/mL of PLGA microparticles.

We next evaluated cellular response to our IMF hydrogels to both validate our DOE results and functionally test the IMF microparticles in 3-dimensions. We investigated whether IMFs could still induce rMSC differentiation and anti-inflammatory macrophage polarization when delivered from microparticles in hydrogels. We loaded 1×10^6^ rMSCs/mL into 10% GelMA with 15 mg/mL IMF microparticles (MP). We noted increased calcium deposition (**Figure 3B**) and AR staining (**Figure 3C**) after 21 days in gels with microparticles and osteogenic media (OM) compared to those treated with OM alone. Impressively, we also observed the greatest increase in ALP activity (**Figure 3A**) in IMF gels with no OM. For macrophage response, we also loaded IMF hydrogels with 1×10^6^ IC-21 cells/mL. After 3 days, we digested the GelMA, retrieved a single cell suspension, and then performed flow cytometry for phenotyping (**Figure 3D**). For this experiment, we included a group with blank MPs to ensure that the acidic byproducts of PLGA degradation were not influencing macrophage polarization [29]. We found no differences in M1 macrophages (**Figure 3E**), but a significant increase in M2 macrophages in our gels with IMF microparticles (**Figure 3F**). For those treated with blank microparticles, we did not detect changes in macrophage phenotype compared to gels without microparticles, indicating that PLGA degradation has no effect on macrophage polarization. Taken together, these results show that the targeted cellular response from our DOE is still maintained with IMFs that are delivered in PLGA microparticles within a hydrogel.

**Figure 3:**
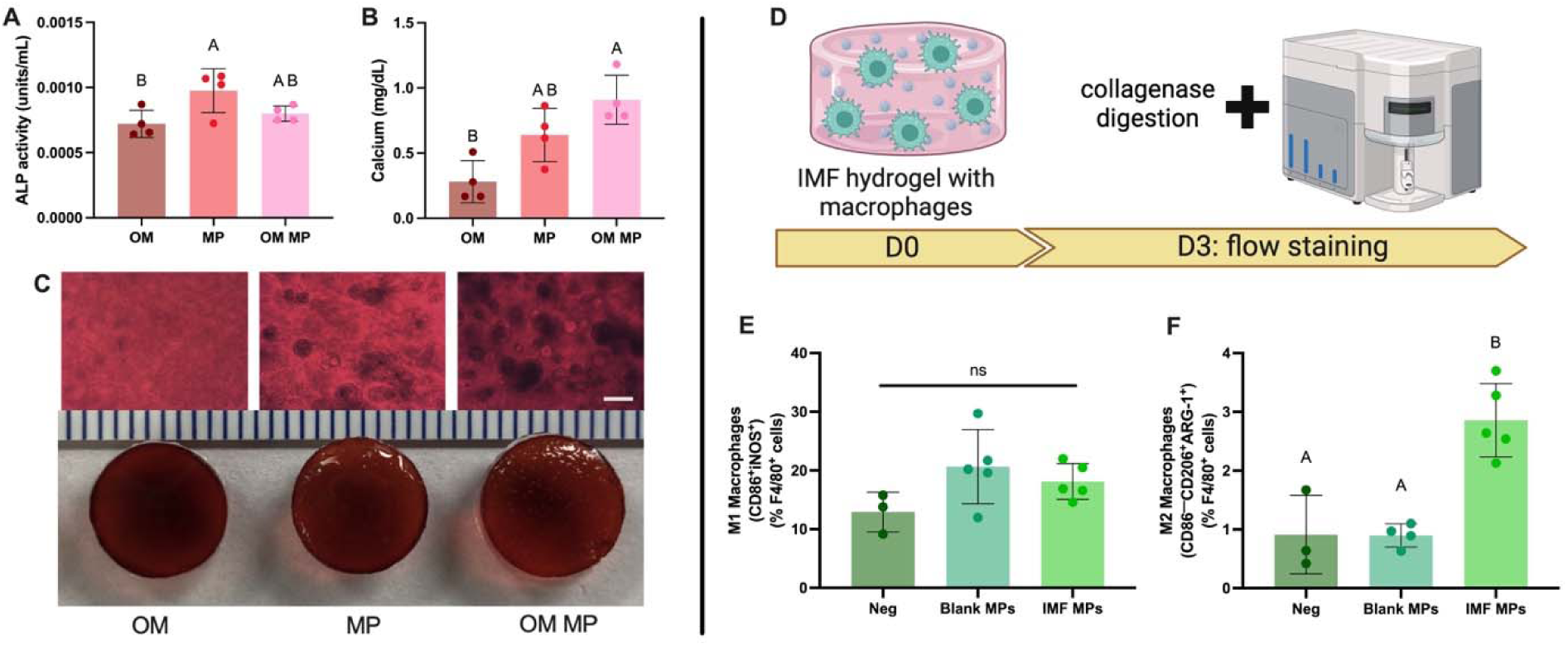
IL-4 and IL-10 delivered in PLGA microparticles within a hydrogel increase rMSC osteogenic differentiation and M2 macrophage polarization. rMSC osteogenic response to IMF hydrogels as measured by **(A)** ALP activity, **(B)** calcium deposition, and **(C)** AR staining after 21 days. Groups include osteogenic media only (OM), IMF microparticles (MP), and IMF microparticles with OM (MP OM). Scale bar on microscopic images = 200 μm. Scale bar on gross images = 1 mm. **(D)** Schematic of workflow of macrophage response to IMF hydrogels. Flow cytometric quantifications of **(E)** CD86^+^iNOS^+^ (M1) macrophages, and **(F)** CD86^-^CD206^+^Arg1^+^ (M2) macrophages. Groups include IC-21s in 10% GelMA only (Neg), with blank microparticles (Blank MPs), and with IMF microparticles (IMF MPs). Data are mean ± SD (n=3-5). Groups with statistically significant differences based on one-way ANOVA do not share the same letters. ns denotes no significance.

To increase the translatability of this model, we interrogated naïve rMSC response under clinically relevant inflammatory conditions. We first characterized how low-level chronic inflammation affects naïve rMSC osteogenic potential. We cultured rMSCs in 10% GelMA hydrogels with basal media supplemented with 50 pg/mL TNFα. We also included monolayer groups to contextualize this work with the literature. After 21 days in culture, we found that *Runx2*, an early osteogenic transcription factor, was significantly upregulated in media with TNFα compared to basal media controls in monolayer and 3D culture (**Figure 4A, 4C,** respectively). *Col1A1*, a late osteogenic marker, had an increasing trend in monolayer between basal media, TNFα supplemented, and TNFα with OM (**Figure 4B**), but only a significant increase in the TNFα OM group in 3D (**Figure 4D**). Together, these results demonstrate that at low levels similar to chronic inflammation, TNFα primes naïve MSCs towards the osteogenic lineage.

**Figure 4:**
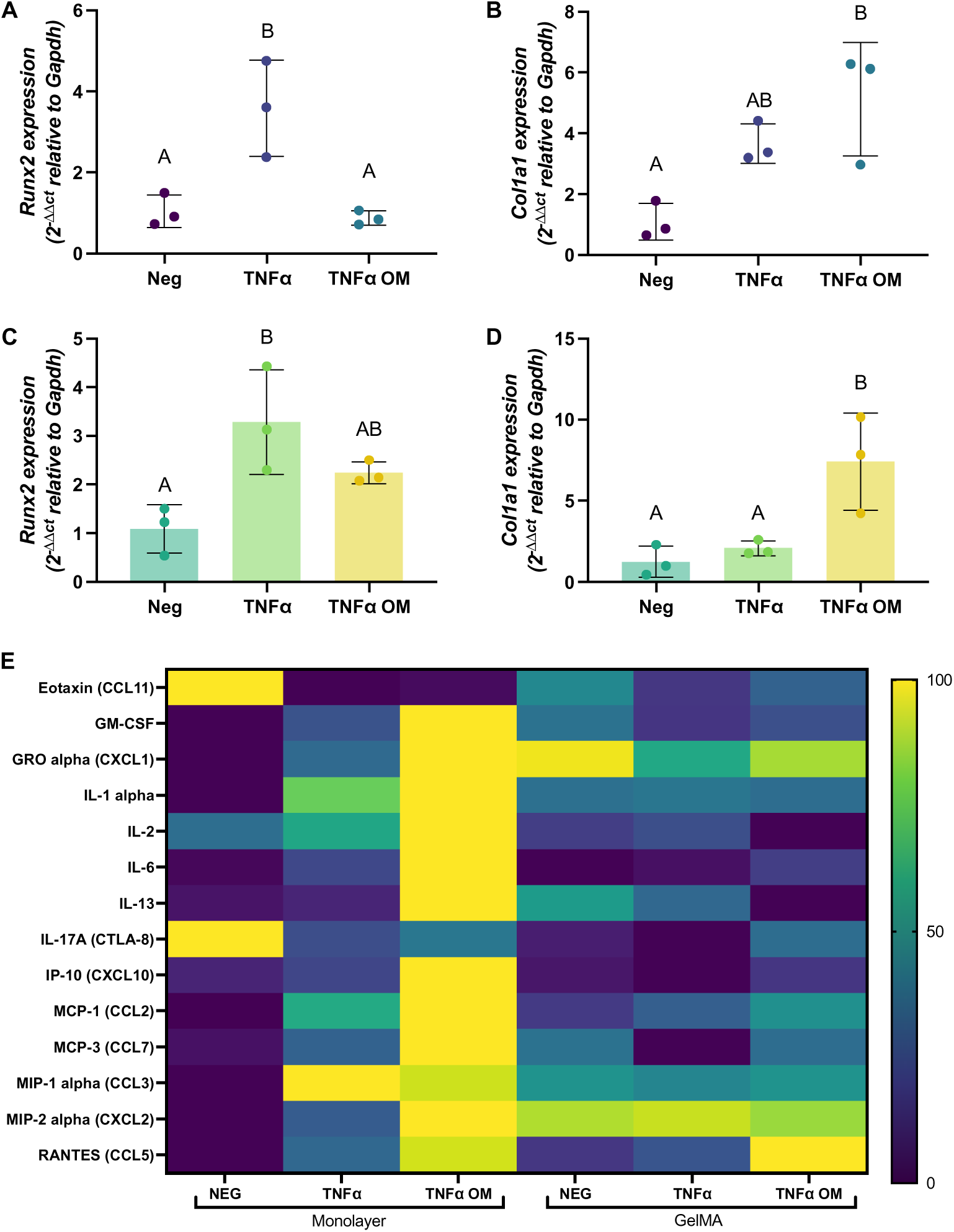
Chronic inflammatory stimulation primes naïve rMSCs towards the osteoblastic lineage. *Runx2* expression at 21 days as an early osteogenic differentiation marker in monolayer **(A)** and 10% GelMA **(C)**. Type 1 collagen (*Col1A1*) expression at 21 days as a marker of late osteogenic differentiation in monolayer **(B)** and 10% GelMA **(D)**. Data are mean ± SD (n=3). Groups with statistically significant differences based on one-way ANOVA do not share the same letters. ns denotes no significance. **(E)** Multiplex proteomic analyses of rMSCs after 21 days; n=3-4, data are presented as mean and are normalized per analyte to a percentile scale where the lowest value is zero and the highest is 100. Groups include basal media (NEG), basal media supplemented with 50 pg/mL TNFα (TNFα) and OM with 50 pg/mL TNFα (TNFα OM).

We also aimed to understand how these TNFα levels may affect the MSC immunomodulatory secretome (**Figure 4E**). rMSCs cultured in monolayer with TNFα and osteogenic media exhibited the greatest upregulation in most factors compared to all other groups. Of note, pro-inflammatory factors IL-1α, IL-2, and IL-6 were most upregulated, along with many of the chemokines, which is congruent with TNFα supplementation and suggests that OM likely augments inflammatory priming. Additionally, secreted factors from rMSCs in 3D culture were predominantly within the range of those in monolayer, implying that 10% GelMA may decrease the influence of TNFα on rMSC secretome response.

Having established that low-level TNFα primes naïve rMSCs towards the osteoblastic lineage, we next tested the ability of our IMF hydrogels under inflammatory conditions to induce differentiation of rMSCs (**Figure 5A**). Briefly, we seeded 1×10^6^ rMSCs/mL in hydrogels without MPs, with blank MPs, and with IMFs, and cultured constructs for 21 days in basal media supplemented with 50 pg/mL TNFα. We then performed qPCR and multiplex proteomic analysis. To elucidate whether supplying exogenous IMFs would decrease rMSC-driven immunomodulation, we also collected conditioned media to evaluate functional immune response with macrophage polarization. We noted significant upregulation in *Runx2* expression in IMF hydrogels (**Figure 5B**). This is noteworthy given that these samples are normalized to the TNFα− only group, which was the most upregulated group in Figure 4 Though not significant, we also observed an increasing trend in *Col1A1* expression (**Figure 5C**). Multiplex proteomics at Day 21 revealed that IMF hydrogels with TNFα produced the highest levels of most of the interrogated cytokines (**Figure 5D**). However, macrophage response to this conditioned media showed no differences in M1 (**Figure 5E**) or M2 (**Figure 5F**) phenotypes between groups of the same timepoint. Interestingly, across all timepoints, we detected no differences at each time point but a general increase in the total percentages of polarized macrophages over time. This could be due to cell proliferation and an associated global increase in secreted factors over time. Of note for this study, the absence of differences in polarized cells at each time point regardless of treatment established that MSCs in IMF hydrogels have no functional differences in secretome production. This finding, in combination with qPCR results, supports our hypothesis that exogenous supplementation of IMFs can indirectly promote MSC osteogenesis.

**Figure 5:**
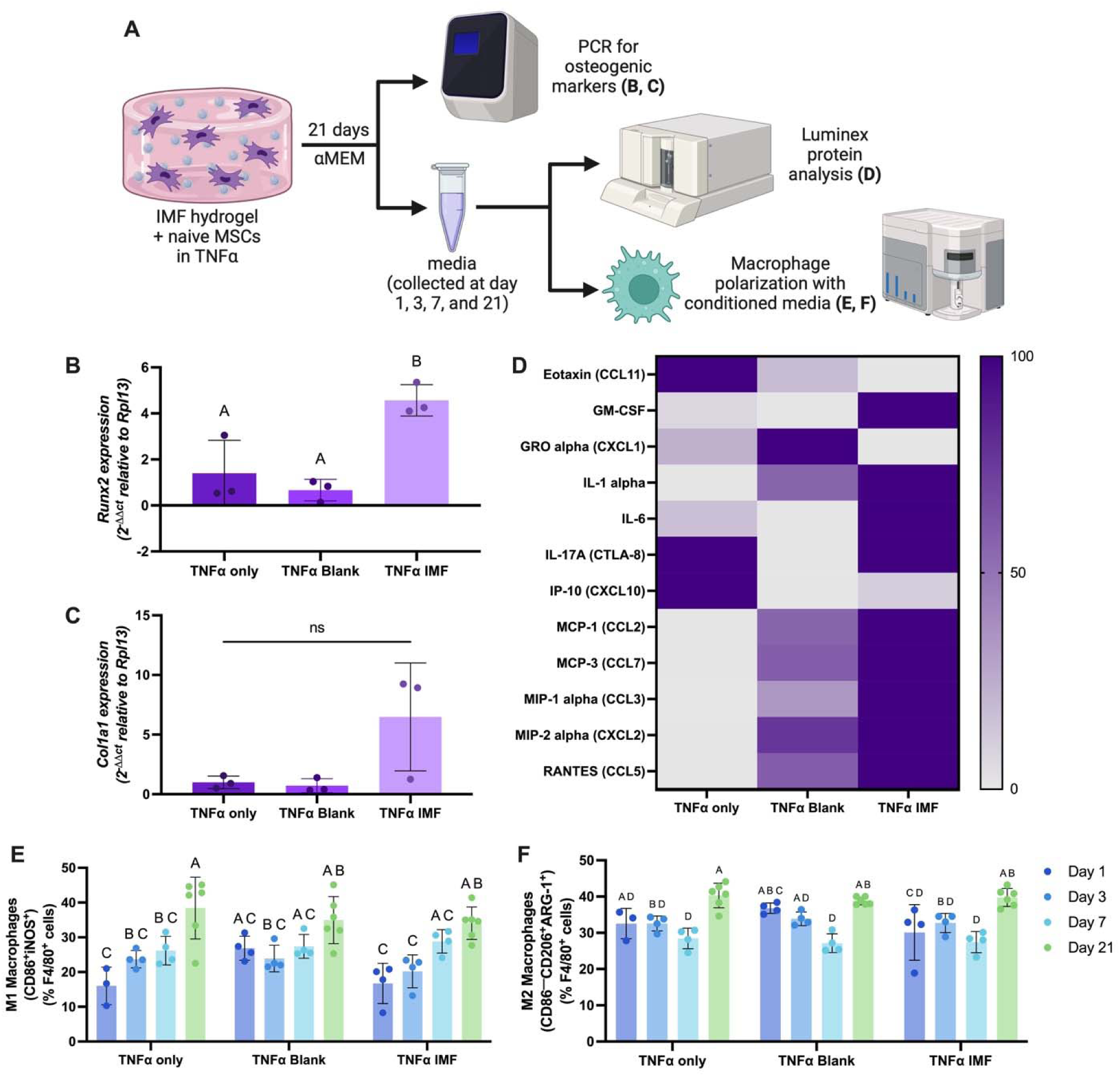
IMF hydrogels in inflammatory conditions promote naïve osteogenic differentiation. **(A)** Schematic of experimental workflow. **(B)** *Runx2* and **(C)** Type 1 collagen (*Col1A1*) expression at 21 days. **(D)** Multiplex proteomic analyses of rMSCs after 21 days; n=5-6, data are presented as mean and are normalized per analyte to a percentile scale where the lowest value is zero and the highest is 100. Flow cytometric quantifications of **(E)** CD86^+^iNOS^+^ (M1) macrophages, and **(F)** CD86^-^CD206^+^Arg1^+^ (M2) macrophages in response to rMSC conditioned media. Data are mean ± SD (n=3-6). Groups with statistically significant differences based on one-way or two-way ANOVA do not share the same letters. ns denotes no significance. Groups are all supplemented with 50 pg/mL TNFα and include basal media (TNFα only), blank MPs (TNFα blank), and IMF microparticles (TNFα IMF).

Having established that IMF hydrogels effectively modulate the immune microenvironment and promote naïve rMSC osteogenic differentiation *in vitro*, we tested the therapeutic potential of naïve MSCs in IMF hydrogels in an *in vivo* model of chronic inflammation: medication-related osteonecrosis of the jaw (MRONJ). MRONJ is a chronic inflammatory disease that results in maxillofacial bone loss and is caused by long-term use of bisphosphonates [39, 40]. The medication of choice to halt bone resorption in osteoporosis and various oncological malignancies [18, 39], bisphosphonates primarily act through systemic high-affinity binding to hydroxyapatite and direct osteoclast inhibition [41]. Unfortunately, MRONJ is a severe complication of this treatment regime, affecting up to 18% of patients taking bisphosphates [42]. If left unchecked, complications can lead to pathological fractures, permanent nerve damage, and complete loss of jaw function [39, 41]. Yet, effective therapies for MRONJ are lacking [42, 43]. Typical treatment options for MRONJ involve surgical debridement, antibiotics, and pain management [18, 42, 43] yet offer only a 50% chance of halting necrosis [44]. Other experimental treatments include low-intensity lasers, platelet rich plasma (PRP), hyperbaric oxygen, and stem cells, all with limited success [42, 43]. This indicates a critical need for novel strategies to support maxillofacial bone health.

Our maxillary model of MRONJ was adapted from previously established models [31], and it was conducted in Sprague Dawley rats as shown in **Figure 6A**. Our study groups included 10% GelMA loaded with rMSCs, IMF microparticles, or both rMSCs and IMF microparticles. Intravascular bisphosphonate injection is well established to create a necrotic, non-healing lesion following tooth extraction [31], so we excluded a true negative control. Twenty-four hours after surgeries and implantation, whole body fluorescent imaging was used as non-invasive surrogate to measure inflammation. Three days and six weeks post-operatively, maxillae were collected to evaluate tissue regeneration, specifically focusing on inflammatory modulation and bone formation, respectively.

**Figure 6:**
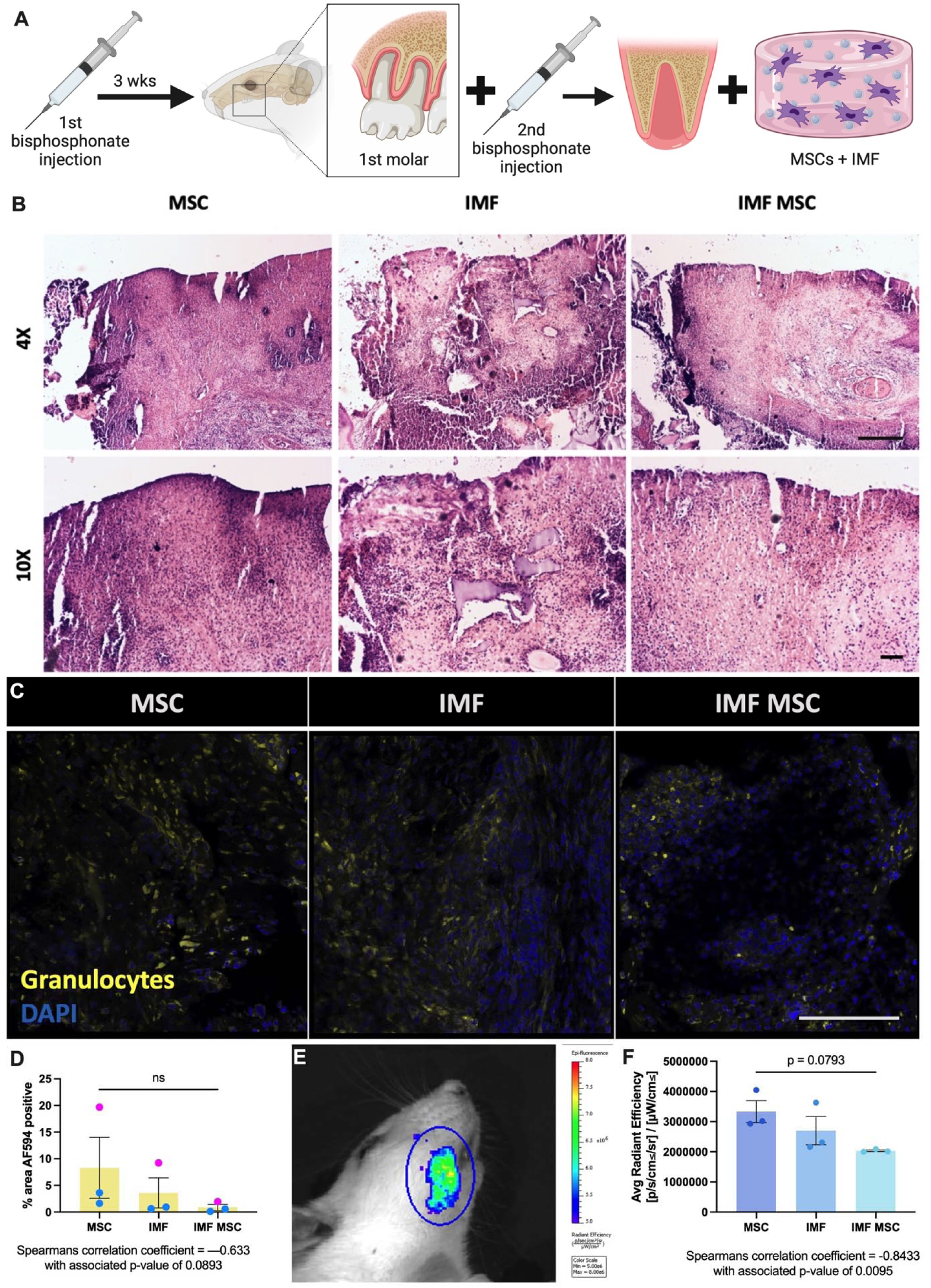
IMF hydrogels with naïve rMSCs decrease early inflammation *in vivo* in MRONJ model. **(A)** Schematic of maxillary MRONJ model. **(B)** Representative images of H&E staining of defect site 3 days after surgery, with low (4X, scale bar = 500 μm, top) and high (10X, scale bar = 100 μm, bottom) magnification views. **(C)** Representative images of immunohistochemical stain for granulocytes in the defect site 3 days after surgery. Scale bar = 100 μm. **(D)** Quantification of granulocyte positive staining, where pink data points are from female specimens and blue are from male. **(E)** Representative image of IVIS scan and ROI with **(F)** quantification of ROIs. Data are mean ± SEM (n=3). p-values shown are based on one-way ANOVA. Below (D) and (F), non-parametric Spearman’s correlation coefficient, r, indicates association; Spearman correlation p-values reports statistical significance of association.

H&E staining of the defect area three days after surgery revealed noticeable hematoxylin-heavy staining in MSC groups that decreased in IMF samples and further decreased in IMF MSC groups (**Figure 6B**), suggesting that inflammatory cell infiltrate decreases across groups. However, immunohistochemical analysis revealed that granulocytes (**Figure 6C**) exhibited a decreasing, though not significant, trend between MSC, IMF, and IMF MSC groups (**Figure 6D**). This corroborates the appearance of the cellular infiltrate noted with H&E, with the likely predominant cell type being neutrophils. Interestingly, female rats appeared to have an approximately 10-fold increase in positive staining, though we did not have enough samples for a sex-based comparison. Analysis with a pan-cathepsin probe to identify localized areas of increased inflammation (**Figure 6E**) revealed a significantly decreasing trend in inflammation across groups 24 hours post-operatively, with a Spearman’s Correlation Coefficient of –0.8433 and associated p-value of 0.0095 (**Figure 6F**). Quantifications are based on male rats only, as female rats had significant background signal, especially around the eye, that overwhelmed the molar ROI. This may again point to inflammatory, sex-based differences. Together, these data show that IMF hydrogels loaded with rMSCs effectively decrease the harsh inflammatory environment *in vivo*.

After six weeks, H&E staining showed healed mucosa and appropriate wound closure in all groups (**Figure 7A**). Of note, IMF MSC stains revealed mucosal morphology with increased definition of the prickle cell layer and an undulating basal cell layer, which is most similar to native oral mucosa [45]. Live animal microcomputed tomography (microCT) scans were taken before tooth extraction (baseline), within 24 hours of surgery (D0), and 6 weeks after surgery (W6). VOIs were drawn plane-by-plane over the extracted molar and associated maxillary bone (**Supplemental Figure S6**) in baseline scans, then D0 and W6 scans were overlaid. Presented bone volume is the change in volume of voxels with hydroxyapatite density greater than 250 mg/mL within the molar VOI between the D0 and W6 scans, thus representing bone regeneration within the molar region. Though there was no difference between groups across all animals (**Figure 7B**), when separated across sexes, female rats showed significant increases in bone volume for MSC and IMF MSC groups (**Figure 7C**). Males showed no differences across all groups (**Figure 7D**). Importantly, there were no decreases in bone volume, indicating that all groups were effective in halting the degenerative process of MRONJ. Mineral densities of the molar and jaw VOIs also showed no differences across all groups (**Figure 7E, 7H**), but again showed significant trends when presented as a function of sex. For the molar, mineral densities in females significantly increased when treated with MSCs; significantly decreased in IMF groups; and did not change with IMF MSC treatment, though the final W6 density between MSC and IMF MSC groups was statistically similar (**Figure 7F**). The males, though not significant, decreased in mineral density across all individuals and samples over time (**Figure 7G**). In the jaw, females showed no significant changes in mineral density, though MSC groups at D0 had significantly lower starting densities at D0 than IMF groups, yet all three treatments had similar densities by W6 (**Figure 7I**). Males again showed general decreases in density, though IMF MSC-treated groups significantly decreased over time (**Figure 7J**). Together, these data show that there are significant differences in regenerative responses to these therapies based on sex and that local delivery of MSCs is crucial for improved bone regeneration.

**Figure 7:**
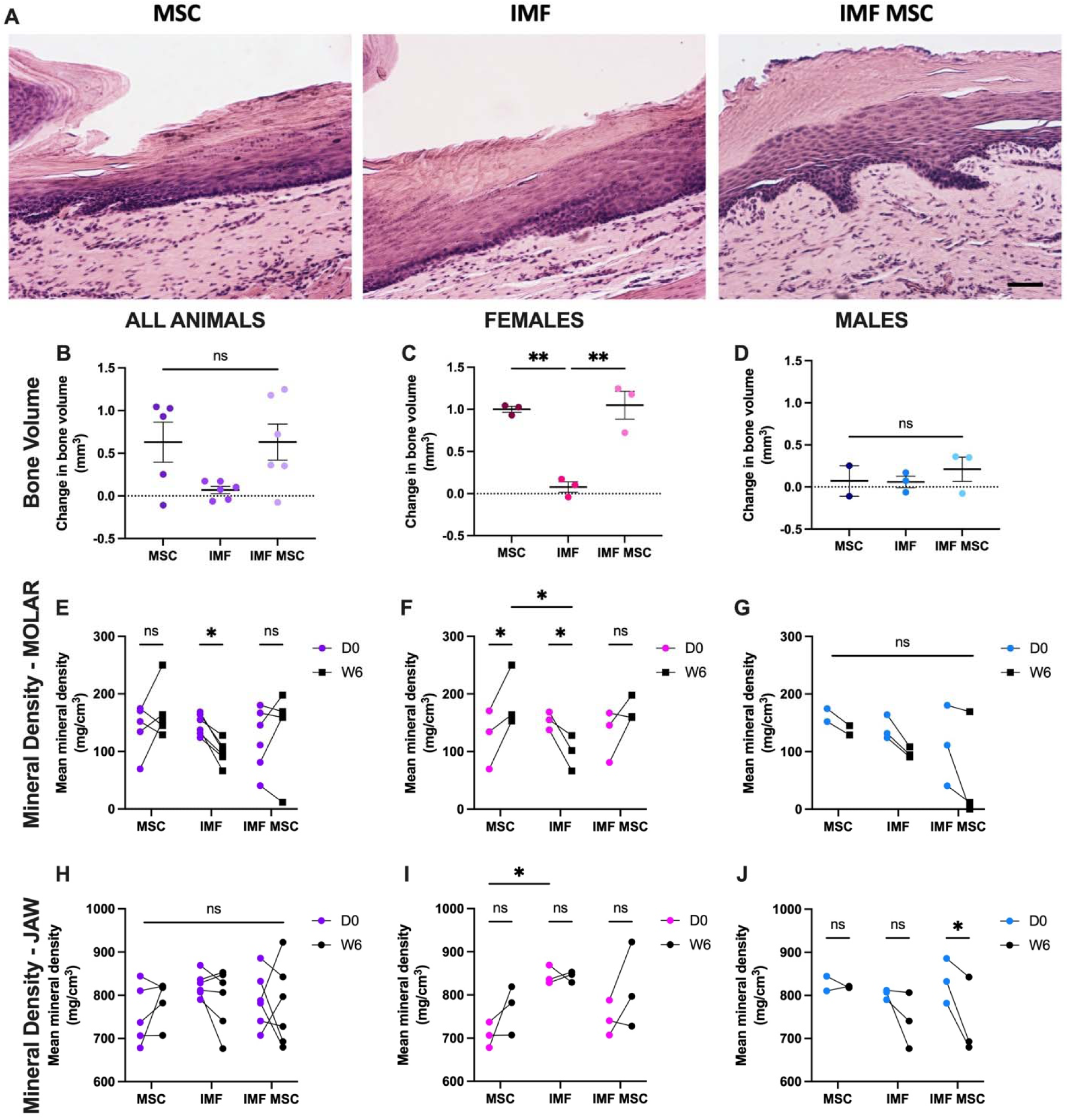
Local delivery of MSCs is crucial for bone regeneration in female rats. **(A)** Representative images of H&E staining of defect site 6 weeks after surgery, scale bar = 100 μm. Quantified change in bone volume within the molar VOI of microCT scans between D0 and W6 of **(B)** all animals, **(C)** females, and **(D)** males. Quantified mineral density of the molar VOI in **(E)** all animals, **(F)** females, and **(G)** males. Quantified mineral density of the maxillary VOI in **(H)** all animals, **(I)** females, and **(J)** males. Data are mean ± SEM (n=5-6 for all animals; n=2-3 for sex differences. Groups with statistically significant differences as determined by one-way or paired two-way ANOVA are denoted as *p < 0.05, **p < 0.01. ns denotes no significance.

Another important feature of MRONJ pathology is significantly decreased angiogenesis, and reintroduction of blood supply is an important factor in estimating treatment success. Within the defect sites, we noted no significant differences in angiogenesis between groups at W6, as indicated by CD31 staining (**Supplemental Figure S7**). However, we observed positive staining throughout our samples, suggesting that blood supply was re-established in all groups and that differences in bone regeneration were, again, likely due to MSC incorporation in our gels.

## 4. Discussion

There is a clinical need to unite tissue differentiation and immune regulation in regenerative medicine to provide sustained healing in chronic disease, especially those associated with dysregulated inflammation. In this work, we report a two-pronged approach with concurrent immune modulation and osteogenesis. Specifically, we designed and fabricated the controlled release of IMFs from a composite microparticle-hydrogel system for localized delivery. The formulation of these IMFs supplied exogenous, pro-regenerative signals to the immune microenvironment, which in turn decreased MSC-mediated immunomodulation and instead promoted differentiation towards the osteoblastic lineage. Utilizing a rat model of MRONJ as a pre-clinical example, we also demonstrate that these techniques are effective *in vivo*. This serves as an important representation of the applicability of this work, as MRONJ is a chronic, debilitating, painful disease with high morbidity rates. Though typical and experimental therapeutics have predominantly focused on halting bone necrosis [18, 42, 43], chronic inflammation and disrupted osteoclastogenesis drive the pathobiology of the disease, suggesting that the limited success of current strategies may be partially due to insufficient consideration of the role of the immune system [46]. Thus, by showing that successful immunomodulation impacts tissue regeneration, this work has the potential to influence therapeutic approaches to chronic inflammatory diseases such as MRONJ.

Inflammatory cues modulate wound healing, including bone regeneration, and MSC osteogenic differentiation requires a temporally and spatially complex mixture of cues from the immune system [6, 32]. For years, local or systemic immunomodulation has been pursued to promote bone healing with varying degrees of success. The canonical example is clinical contraindications for non-steroidal anti-inflammatory drugs (NSAID) in fracture patients [47]. Previously prescribed for analgesia, there is now a significant collection of literature showing that systemic and local NSAID use prolongs healing time and increases the risk of non-unions, likely due to disrupted TGF-β3 signaling during endochondral ossification [48]. Specific immunomodulatory drugs [49], such as prostacyclin [50] or Myd88 antagonists [51], can improve poor clinical outcomes, such as non-unions or fractures in the elderly, both of which are known to have dysregulated inflammatory responses that precipitate poor regeneration. However, these treatments are often injected systemically or locally as a liquid solution, thus failing to provide temporal and spatial instruction. Here, we reported that localized, controlled delivery of IMFs modulates immune response, specifically through macrophage polarization, and improves osteogenesis. Growth factor delivery *via* polymeric microparticles is well characterized for a variety of applications, from chemotherapeutic delivery [52] to immune cell instruction [28, 53] to sustained antibiotic release [54]. Of significant relevance, a few studies induced regulatory T cell differentiation in a periodontitis model, showing immunomodulatory responses that secondarily decreased alveolar bone resorption in mice and dogs [55, 56]. However, the innovation of our work lies largely in its application towards MRONJ, localized delivery within a hydrogel, and rigorous approach to identify an optimal IMF formula for simultaneous immunomodulation and bone formation. To our knowledge, this is the first study to report such an experimental design with *in vitro* and *in vivo* analyses, all with an eye towards translational implementation.

We also show significantly different responses to IMF treatment *in vivo* as a function of sex. Over 80% of MRONJ patients are female due to the high correlation of MRONJ as a complication of osteoporosis and breast cancer treatments [40]. However, sex differences persist in clinical bone healing and inflammation beyond MRONJ. In general, female fracture healing time is extended compared to males, and they are at increased risk for atrophic non-unions [57, 58]. Hormonal differences represent obvious culprits for these discrepancies [57, 58], but general immune and inflammatory profiles that differ between sexes [59] may be another potential element. In fact, females account for over 80% of patients with autoimmune disorders [60], indicating inherent differences in baseline immune systems. Indeed, inflammatory Type I and Type II interferon signaling is increased in females across numerous species [60]. Thus, taken together, it is conceivable that females may have improved or exaggerated responses to immunomodulation compared to males, as we see here. Interestingly, sex differences in MRONJ treatment studies are often not reported, highlighting another novel component to this work.

Despite the innovation and interesting findings presented herein, this work has several limitations. Microbial elements remain unexplored and are a major confounding factor. Clinical MRONJ therapies include strong topical and systemic antimicrobial treatments to help resolve associated inflammation [43]. We neither treated with antibiotics nor analyzed changes in the oral microbiome. Given the immunomodulatory mechanism of our IMF hydrogels, it will be critical to understand the interplay between inflammatory downregulation and opportunistic infections. However, avoiding treatment with antimicrobials also represents a conservative experimental design, suggesting that tissue healing occurred despite the probable presence of opportunistic bacteria. Thus, it is likely that under clinically relevant antimicrobial therapy, there may be improved tissue regeneration due to the reduction of oral microbes and associated immune reactions. We also did not test these constructs retroactively once necrosis had begun. While our IMF hydrogels could conceivably function proactively at the time of dental procedures, as spontaneous occurrence of MRONJ accounts for less than 15% of cases [40], it is an important consideration for future work. Lastly, we did not interrogate different release kinetics of our IMF hydrogels, which may provide clinical flexibility in application and efficacy. Future studies are needed to incorporate these various components.

## Conclusion

These studies report localized, controlled delivery of IMFs to simultaneously modulate the inflammatory environment and improve osteogenesis. These data represent key findings that could facilitate improved treatment of chronic, inflammatory bone loss, such as occurs in MRONJ, as well as other applicable conditions such as nonunion fractures, osteomyelitis, and rheumatic conditions.

## Data availability statement

The data that support the findings of this study are available from the corresponding author upon reasonable request.

## Acknowledgements

Research reported in this publication was supported by the National Institutes of Health under award numbers R01 DE025899 and R01 AR079211 to JKL. The content is solely the responsibility of the authors and does not necessarily represent the official views of the National Institutes of Health. The funders had no role in the decision to publish, or preparation of the manuscript. This work was supported in part by the UC Davis School of Veterinary Medicine Endowment Funds and Graduate Student Support Program to KHG. JKL also acknowledges financial support from the Lawrence J. Ellison Endowed Chair of Musculoskeletal Research. The authors would also like to thank Allen Tu for assistance in the synthesis and characterization of the IMF microparticles. BioRender was used to make all schematics.

## Supplemental Information

**Supplemental Figure S1:**
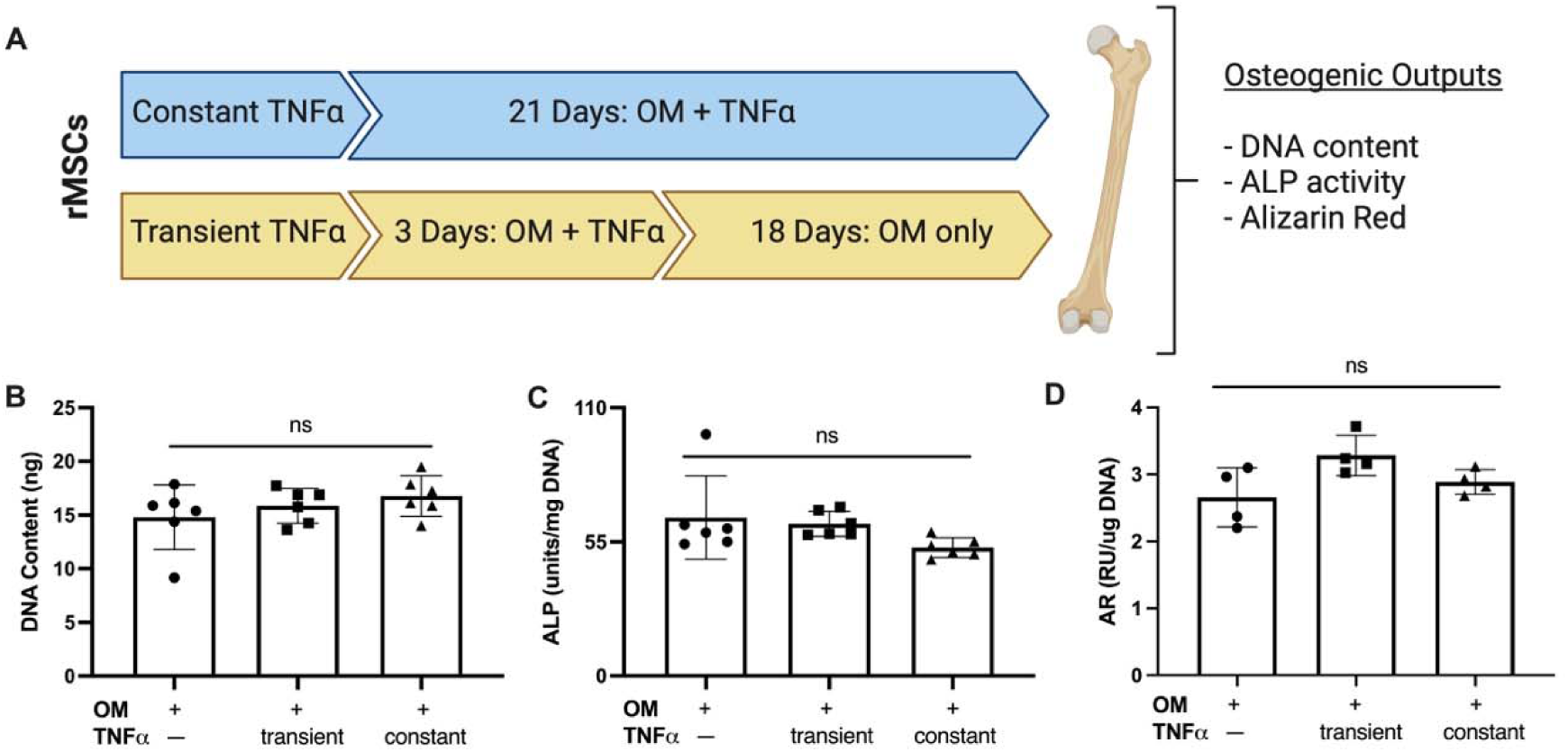
rMSC osteogenesis is not inhibited when treated with 50 pg/mL TNFα. **(A)** Schematic of study timeline and methods. **(B)** DNA content, **(C)** alkaline phosphatase (ALP) activity and **(D)** Alizarin Red (AR) quantifications of rMSCs in monolayer culture after 21 days. Data are mean ± SD (n=4-6). ns denotes no significance based on on-way ANOVA.

**Supplemental Table 1:** DOE experimental regime for macrophage polarization experiment.

| Std | Run | Factor 1 | Factor 2 | Factor 3 | Per 4 mL |  |  |
| --- | --- | --- | --- | --- | --- | --- | --- |
|  |  | A:IL-10.con<br>ng/mL | B:IL-4.con<br>ng/mL | C:PGE2.con<br>ug/mL | IL-10 VOLUME | IL-4 VOLUME | PGE2 VOLUME |
| 8 | 1 | 100 | 50 | 2 | 4 uL | 2 uL | 8 uL |
| 12 | 2 | 50 | 100 | 2 | 2 | 4 | 8 |
| 14 | 3 | 50 | 50 | 1 | 2 | 2 | 4 |
| 9 | 4 | 50 | 0 | 0 | 2 | 0 | 0 |
| 15 | 5 | 50 | 50 | 1 | 2 | 2 | 4 |
| 2 | 6 | 100 | 0 | 1 | 4 | 0 | 4 |
| 10 | 7 | 50 | 100 | 0 | 2 | 4 | 0 |
| 16 | 8 | 50 | 50 | 1 | 2 | 2 | 4 |
| 1 | 9 | 0 | 0 | 1 | 0 | 0 | 4 |
| 13 | 10 | 50 | 50 | 1 | 2 | 2 | 4 |
| 17 | 11 | 50 | 50 | 1 | 2 | 2 | 4 |
| 4 | 12 | 100 | 100 | 1 | 4 | 4 | 4 |
| 11 | 13 | 50 | 0 | 2 | 2 | 0 | 8 |
| 7 | 14 | 0 | 50 | 2 | 0 | 2 | 8 |
| 3 | 15 | 0 | 100 | 1 | 0 | 4 | 4 |
| 5 | 16 | 0 | 50 | 0 | 0 | 2 | 0 |
| 6 | 17 | 100 | 50 | 0 | 4 | 2 | 0 |

**Supplemental Table 2:** DOE experimental regime for rMSC osteogenic experiments.

| Std | Run | Factor 1 | Factor 2 | Factor 3 | Per 5 mL |  |  |
| --- | --- | --- | --- | --- | --- | --- | --- |
|  |  | A:IL-10.con<br>ng/mL | B:IL-4.con<br>ng/mL | C:PGE2.con<br>ug/mL | IL-10 VOLUME | IL-4 VOLUME | PGE2 VOLUME |
| 8 | 1 | 100 | 50 | 2 | 5 uL | 2.5 uL | 10 uL |
| 12 | 2 | 50 | 100 | 2 | 2.5 | 5 | 10 |
| 14 | 3 | 50 | 50 | 1 | 2.5 | 2.5 | 5 |
| 9 | 4 | 50 | 0 | 0 | 2.5 | 0 | 0 |
| 15 | 5 | 50 | 50 | 1 | 2.5 | 2.5 | 5 |
| 2 | 6 | 100 | 0 | 1 | 5 | 0 | 5 |
| 10 | 7 | 50 | 100 | 0 | 2.5 | 5 | 0 |
| 16 | 8 | 50 | 50 | 1 | 2.5 | 2.5 | 5 |
| 1 | 9 | 0 | 0 | 1 | 0 | 0 | 5 |
| 13 | 10 | 50 | 50 | 1 | 2.5 | 2.5 | 5 |
| 17 | 11 | 50 | 50 | 1 | 2.5 | 2.5 | 5 |
| 4 | 12 | 100 | 100 | 1 | 5 | 5 | 5 |
| 11 | 13 | 50 | 0 | 2 | 2.5 | 0 | 10 |
| 7 | 14 | 0 | 50 | 2 | 0 | 2.5 | 10 |
| 3 | 15 | 0 | 100 | 1 | 0 | 5 | 5 |
| 5 | 16 | 0 | 50 | 0 | 0 | 2.5 | 0 |
| 6 | 17 | 100 | 50 | 0 | 5 | 2.5 | 0 |

**Supplemental Figure S2:**
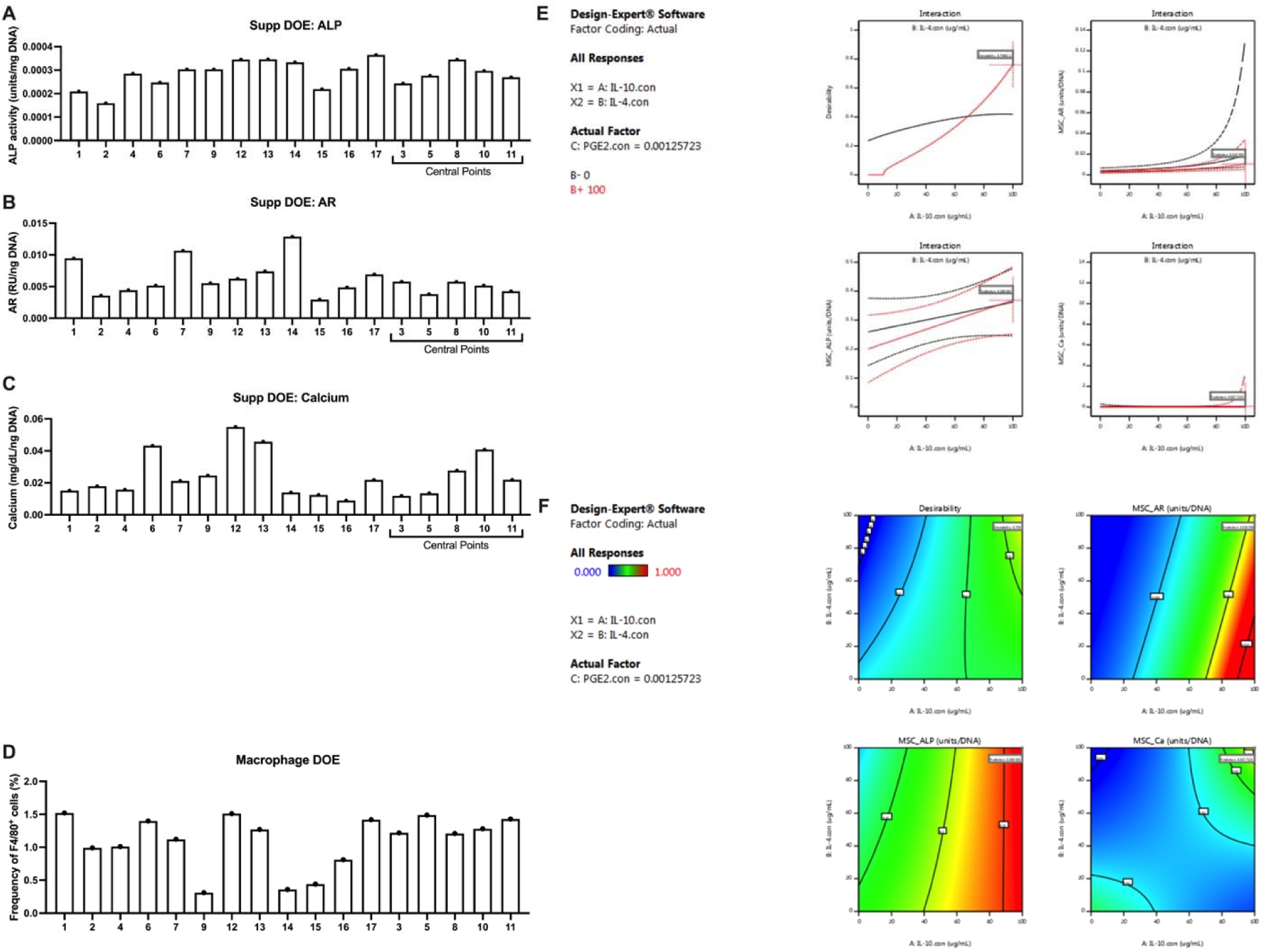
Direct DOE assay results and additional software outputs. Osteogenic results: **(A)** ALP activity, **(B)** AR quantification, **(C)** calcium deposition. **(D)** Macrophage polarization results. **(E)** Predicted interaction plots for IL-4, IL-10, and PGE_2_. **(F)** Additional surface response graphs for all osteogenic results. Data are mean (n=1 per Box Behken Design).

**Supplemental Figure S3:**
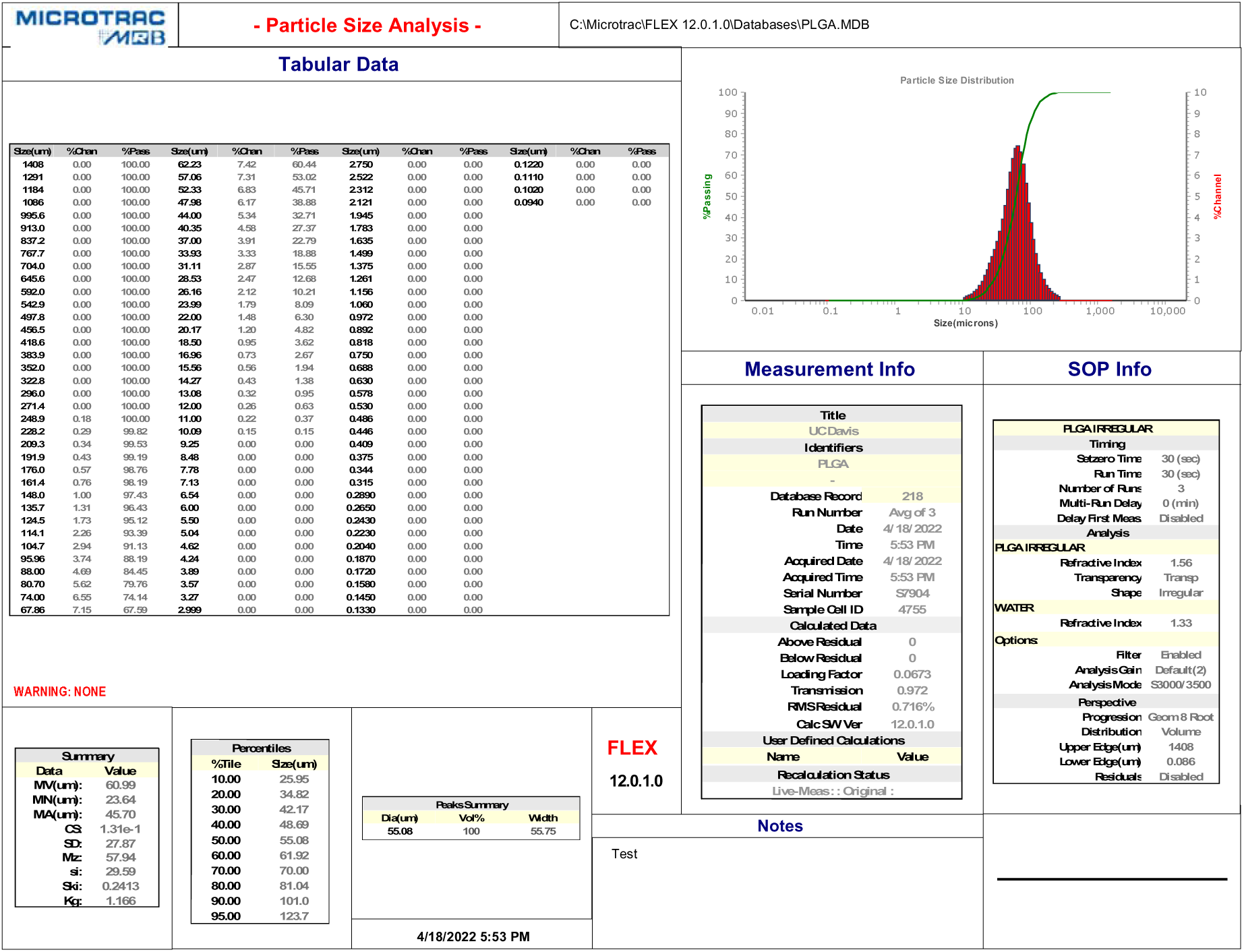
Generated distribution data from S3500 Laser Diffraction Analyzer for IL-4-loaded microparticles.

**Supplemental Figure S4:**
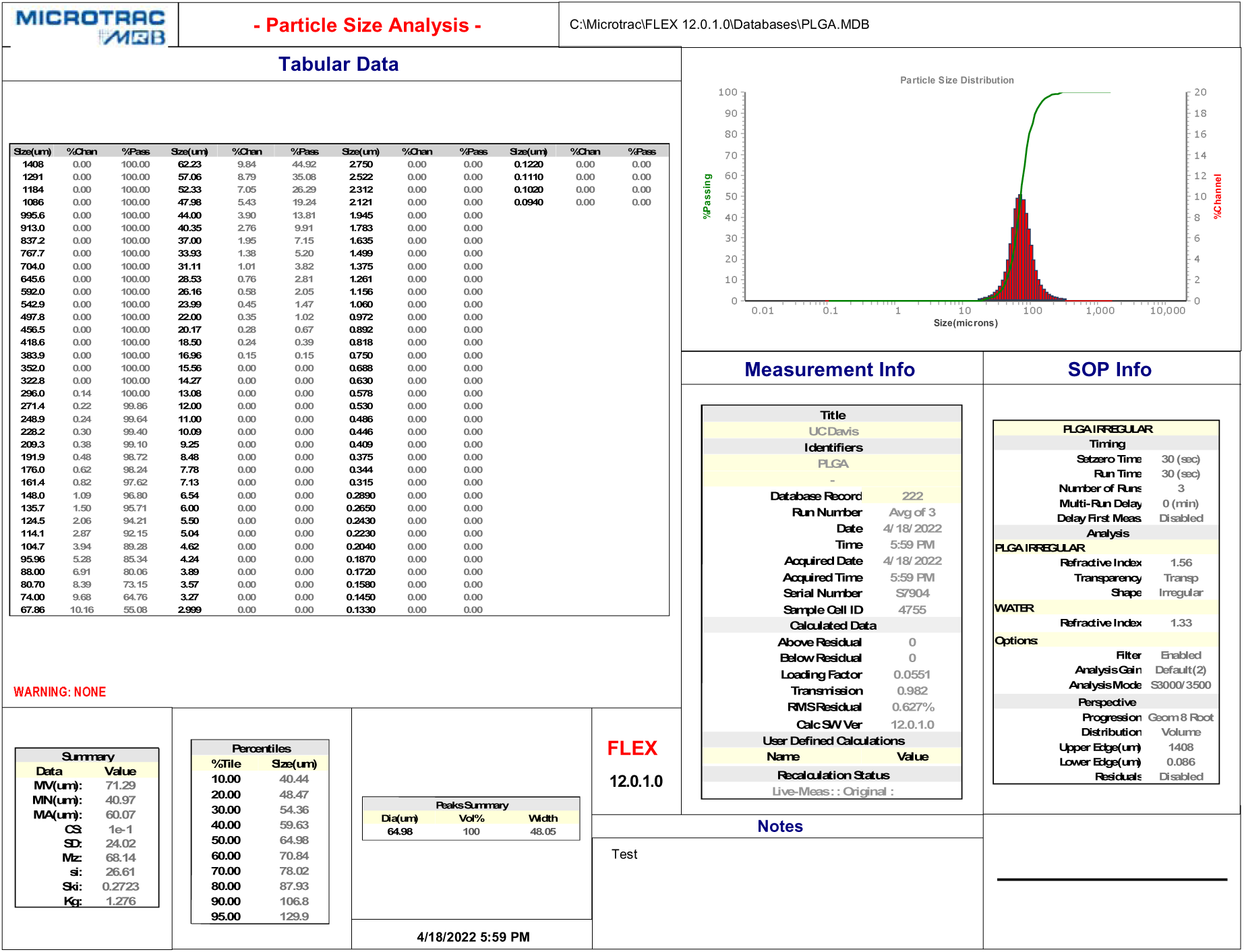
Generated distribution data from S3500 Laser Diffraction Analyzer for IL-10-loaded microparticles.

**Supplemental Figure S5:**
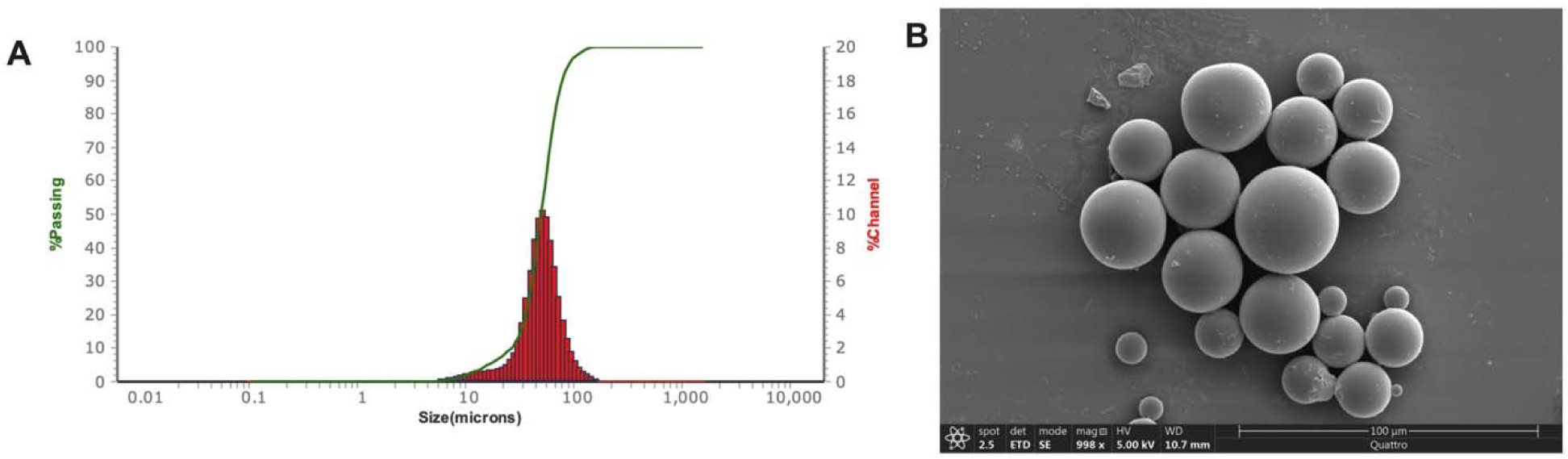
Blank PLGA microparticle characterization. **(A)** Size distribution graph and **(B)** representative SEM image of blank microparticles. Scale bar = 100 μm

**Supplemental Figure S6:**
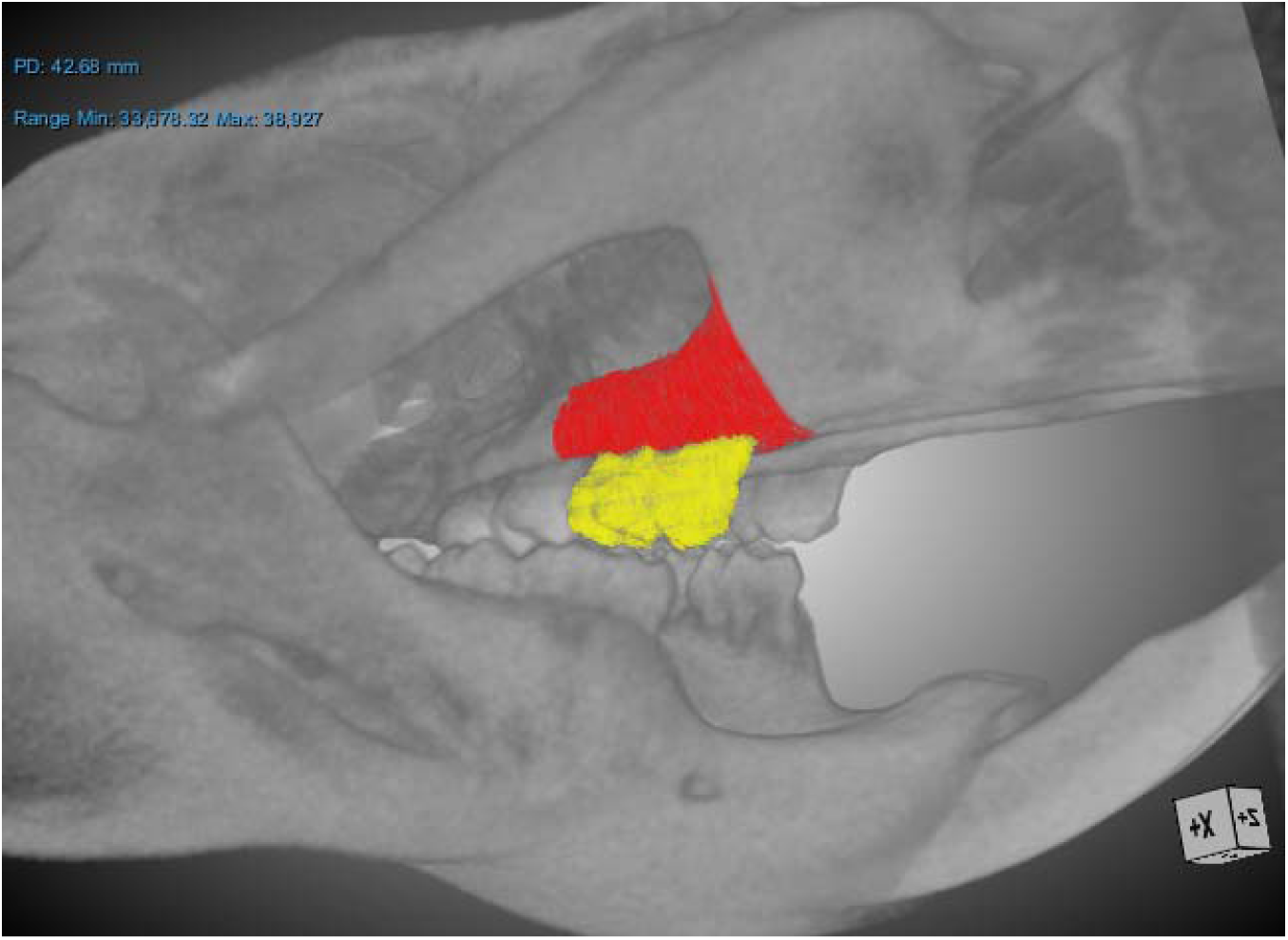
Representative 3D rendering of microCT with selected Molar (yellow) and Jaw (red) VOIs.

**Supplemental Figure S7:**
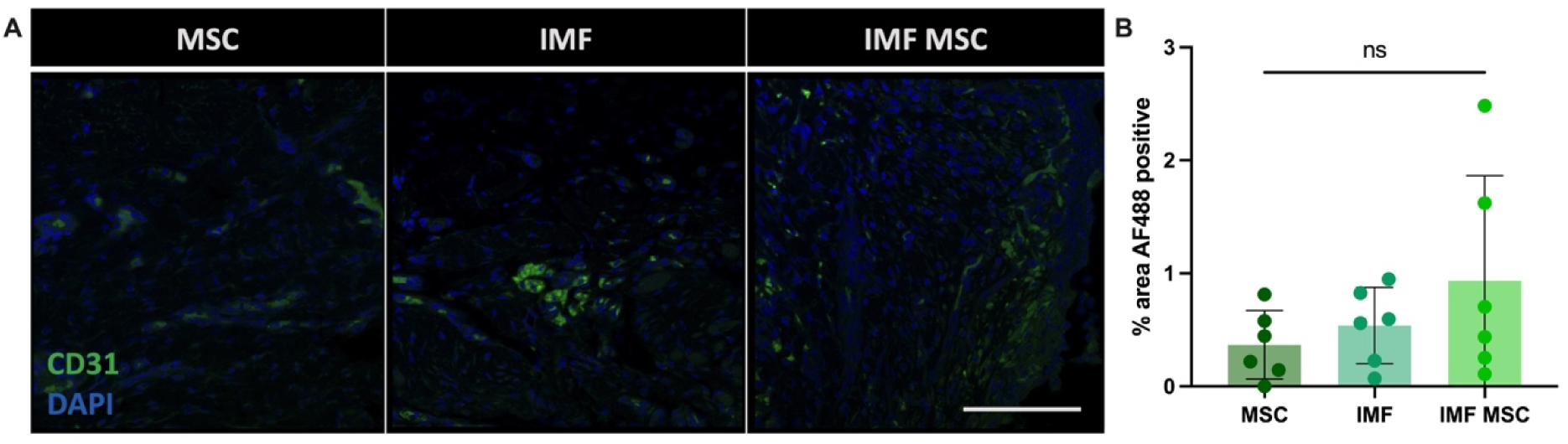
CD31 staining at W6. **(A)** Representative images of CD31 immunohistochemical stain for endothelial cells in the defect site 6 weeks after surgery. Scale bar = 100 μm. **(B)** Quantification of CD31 positive staining

